# A genetic screen for genes that impact peroxisomes in *Drosophila* identifies candidate genes for human disease

**DOI:** 10.1101/704122

**Authors:** Hillary K. Graves, Sharayu Jangam, Kai Li Tan, Antonella Pignata, Elaine S. Seto, Shinya Yamamoto, Michael F. Wangler

**Affiliations:** Department of Molecular and Human Genetics, Baylor College of Medicine (BCM), Houston, TX 77030, USA; Department of Neuroscience, BCM, Houston, TX 77030, USA; Program in Developmental Biology, BCM, Houston, TX 77030, USA; Jan and Dan Duncan Neurological Research Institute, Texas Children Hospital, Houston, TX 77030, USA

**Keywords:** *Drosophila*, peroxisomes, *BRD4*, *fs(1)h*

## Abstract

Peroxisomes are sub-cellular organelles that are essential for proper function of eukaryotic cells. In addition to being the sites of a variety of oxidative reactions, they are crucial regulators of lipid metabolism. Peroxisome loss or dysfunction leads to multisystem diseases in humans that strongly affects the nervous system. In order to uncover previously unidentified genes and mechanisms that impact peroxisomes, we conducted a genetic screen on a collection of lethal mutations on the *X* chromosome in *Drosophila*. Using the number, size and morphology of GFP tagged peroxisomes as a readout, we screened for mutations that altered the number and morphology of peroxisomes based on clonal analysis and confocal microscopy. From this screen, we identified 18 genes that cause increases in peroxisome number or altered morphology when mutated. We examined the human homologs of these genes and found that they are involved in a diverse array of cellular processes. Interestingly, the human homologs from the *X*-chromosome collection are under selective constraint in human populations and are good candidate genes particularly for dominant genetic disease. This *in vivo* screening approach for peroxisome defects allows identification of novel genes that impact peroxisomes *in vivo* in a multicellular organism and is a valuable platform to discover genes potentially involved in dominant disease that could affect peroxisomes.

## Introduction

Peroxisomes are sub-cellular organelles that mediate crucial biological processes in eukaryotic cells, including oxidative reactions, catabolism of very-long-chain fatty acids, catabolism of branched chain fatty acids, catabolism of bile acids, and biosynthesis of plasmalogen lipids (Fujiki *et al*. 2014; Wanders 2014). Human diseases due to lack of peroxisomes are devastating multisystem diseases that result in severe brain, liver, bone and kidney disease (Wanders and Waterham 2005). These conditions, called peroxisome biogenesis disorders Zellweger-spectrum disorders (PBD-ZSD), are a group of multi-system autosomal recessive disorders with severe central nervous system (CNS) manifestations and as yet no effective treatments exists for the hearing, visual and CNS phenotypes (Klouwer *et al*. 2015; Braverman *et al*. 2016).

Historically, genetic screens for biochemical phenotypes have identified genes implicated in peroxisome-biogenesis, pexophagy, and peroxisomal biochemistry (Subramani 1998; Kao *et al*. 2018). More recently, microscopy-based screens have uncovered genes implicated in peroxisome morphology (Baron *et al*. 2016; Yofe *et al*. 2017). These studies have shown that the pathways that regulate peroxisome dynamics (i.e. peroxisome size and number) remain incompletely understood (Mast *et al*. 2015). This is especially true with regards to information gained from multicellular organisms as opposed to yeast and cell models because most essential genes are not amenable to peroxisomal studies due to early lethality in development. Although null alleles in some of the *pex* genes in the fruit fly *Drosophila melanogaster* produce viable adult flies that that exhibit locomotor defects and reduced longevity (Nakayama *et al*. 2011; Faust *et al*. 2014; Wangler *et al*. 2017a), a number of essential genes that regulate peroxisome biogenesis and homeostasis may remain unstudied. To uncover unidentified mechanisms that impact peroxisome number and size, we conducted a forward genetic screen on the *X* chromosome in *Drosophila*. We had previously generated a large collection of recessive lethal mutant lines on an isogenized chromosome and these lines have been extensively screened for developmental and neurological phenotypes (Yamamoto *et al*. 2014). Moreover, we previously utilized this collection of mutations to uncover new human disease genes and showed that human orthologs of these fly essential genes are enriched for genes listed in the Online Mendelian Inheritance in Man (OMIM) disease database (Yamamoto *et al*. 2014; Yoon *et al*. 2017; Tan *et al*. 2018). We took 215 lines from this collection that correspond to 100 genes (98 mapped genes and 2 unmapped complementation group) and screened for peroxisomal phenotypes using GFP tagged peroxisomes (Chao *et al*. 2016; Wangler *et al*. 2017a) in conjunction with clonal analysis, allowing generation of homozygous mutant cells within *Drosophila* larval fat body in an otherwise heterozygous animal to bypass early lethality.

Our screen identified a number of genes not previously implicated in peroxisome dynamics or regulation. In previous studies, we’ve shown that the genes from this collection are enriched for human disease genes (Yamamoto *et al*. 2014). Based on this we propose our screen results as identifying candidate human disease genes particularly for dominant disease that may impact peroxisomes.

## Methods & Materials

### Drosophila X-Chromosome Peroxisome (X-Pex) screen

All *X*-linked recessive lethal mutant alleles utilized in this paper listed in Table 1 and Supplemental Table 1 were generated on an isogenized *y^1^ w^*^ FRT19A* chromosome using ethyl methanesulfonate (EMS) mutagenesis as described (Haelterman *et al*. 2014; Yamamoto *et al*. 2014; Deal and Yamamoto 2018). These fly strains are publically available from the Kyoto Stock Center (https://kyotofly.kit.jp/stocks/documents/EMS_X_lethals.html) or the Bloomington Drosophila Stock Center (https://bdsc.indiana.edu/stocks/chemically_induced_mutations/xlethals.html).

**Table 1.** Hits from the Peroxisome X-pex screen. The Fly gene and specific allele are listed along with the phenotype observed in the screen (Category A, B and C). The human homologs of each gene were identified using DIOPT or HCOP (Hu *et al*. 2011; Braschi *et al*. 2019). Known biological function of the fly protein is listed according to the annotation in UniProt (Uniprot 2019).

| Fly Gene | Allele (s) | Peroxisomal Phenotype | Human Gene(s) | Biological Function in Fly (UniProt) |
| --- | --- | --- | --- | --- |
| *fs(1)h* | *Fs[1]h[A], [B], [C]* | Category A & B | BRD2, BRDT, BRD3, BRD4 | Transcriptional regulation |
| *Rbcn-3B* | *Rbcn-3B[A], [B]* | Category A | WDR7 | Vacuolar acidification, Notch signaling |
| *Coq8* | *Coq8[A],[B]* | Category A | COQ8B | Protein kinase, electron transport |
| *Usp16-45* | *Usp16-45[A],[B]* | Category A | USP45 | Protein deubiquitination |
| *mxc* | *mxc[A],[B],[C],[D], [E]* | Category A | No Human Ortholog | Transcriptional, hemocyte differentiation and proliferation |
| *Cp7Fb* | *Cp7Fb[B]* | Category A | No Human Ortholog | Chorion |
| *Upf1* | *Upf1[A],[B]* | Category A | UPF1 | Nonsense mediated decay |
| *Upf2* | *Upf2[A],[B]* | Category A | UPF2 | Nonsense mediated decay |
| *Nrg* | *Nrg[XB]* | Category A | NRCAM, NFASC, L1CAM, CHL1 | Cell adhesion |
| *Fum1* | *Fum1[A]* | Category A | FH | TCA cycle enzyme (mitochondria) |
| *Coq7* | *Coq7[B]* | Category A | COQ7 | Ubiquinone biosynthesis (mitochondria) |
| *sgg* | *sgg[A],[B],[E]* | Category A | GSK3B, GSK3A | Protein kinase |
| *CG17829* | *CG17829[A],[B]* | Category A | HINFP | Transcriptional regulation |
| *CG3149* | *CG3149[B]* | Category A | RFT1 | Glycolipid translocation |
| *Smox* | *Smox[B]* | Category A | SMAD3 | Transcriptional regulation |
| *PI4KIIIa* | *PI4KIIIa [E], [W]* | Category A | PI4KA | Synaptic growth, cell polarity, membrane organization |
| *MTPAP* | *MTPAP[A],[B]* | Category A | MTPAP | Mitochondrial transcription |
| *temp* | *Temp [A], [B] & [D]* | Category A | PTAR1 | Rab geranylgeranyltransferase activity |
| **TOTAL** | **37 hits** | **18 fly genes** | **23 human genes** |  |

Heterozygous females (*y^1^ w^*^ mut* FRT19A/FM7c Kr-GAL4, UAS-GFP*, mut* indicates the mutation of interest) from these lethal lines were crossed to males of the genotype *hsFLP, Ubi-RFP FRT19A; Actin-GAL4, UAS-GFP-SKL/CyO*, and their embryonic progeny were heat shocked at 0-4 hours after egg laying at 37°C for 1 hour. Third larval instar wandering larvae were dissected in PBS and fixed in 4% paraformaldehyde for 20-30 minutes. Fat bodies were mounted in DAPI (4’,6-diamidino-2-phenylindole) containing mounting media (Vectashield), confocal microscopy images were captured on a LSM 710 laser scanning confocal microscope (Zeiss), and processed in Photoshop (Adobe). The fat body experiments were a peroxisome focused secondary screen on a subset of mutants of the larger X-lethal collection reported in Yamamoto et al. (2014), similar to an Atg8 and *LAMP1* based screen to identify novel autophagy regulators conducted on the same collection (Fang *et al*. 2016). Similar secondary screen were performed on the same X-lethal collection to identify regulators of other biological processes such as ring canal formation and somatic stem cell maintenance during oogenesis (Yamamoto *et al*. 2013; Cook *et al*. 2017) demonstrating the value of this collection in screening for genes involved in diverse cellular processes.

### Human gene candidate analysis

Human homologs from the fly genes were determined using the Human Gene Nomenclature Orthology Prediction (HCOP, https://www.genenames.org/tools/hcop/) and the Drosophila RNAi Screening center Integrative Ortholog Prediction Tool (DIOPT, https://www.flyrnai.org/cgi-bin/DRSC_orthologs.pl) tools (Hu *et al*. 2011; Gray *et al*. 2015). These genes were further examined in a series of public human and model organism databases using the MARRVEL tool (http://marrvel.org/) to gather information about the homologous proteins in human and other model organisms (Wang *et al*. 2017). Human gene nomenclature was confirmed using the HGNC (HUGO Gene Nomenclature Committee, https://www.genenames.org) database (Braschi *et al*. 2019). Mendelian disease links were explored in the OMIM (https://www.omim.org/) database (Amberger *et al*. 2015), and each gene was examined using the gnomAD (https://gnomad.broadinstitute.org/) browser (Lek *et al*. 2016). Each gene was also examined using the DOMINO tool for predicted likelihood of a gene having dominant impact on disease (Quinodoz *et al*. 2017). In addition, *de novo* events were examined in denovo-db (http://denovo-db.gs.washington.edu/denovo-db/) website (Turner *et al*. 2017).

### Data and Reagent Availability

All the supplemental data files are available on the GSA Figshare portal including the Supplemental Tables listed in the manuscript. Supplemental Table 1 lists all the Drosophila reagents that are available from the X-screen through public stock centers including the Bloomington stock center and Kyoto Stock center.

## Results and Discussion

### Identification of genes involved in peroxisomal dynamics

To identify new regulators of peroxisomal morphology and dynamics, we performed a secondary screen on a collection of recessive lethal X-chromosome mutants that are enriched for genes that are homologs to human disease genes (Yamamoto *et al*. 2014; Deal and Yamamoto 2018). It is important to note that X-chromosome contains ~15% of protein coding genes in *Drosophila*, and there are no correlation between X-linked genes in flies and humans. In this screen, we created homozygous mutant clones in the fat body of developing *Drosophila* larvae in an otherwise heterozygous animal and assayed for changes in the distribution pattern of a peroxisomal reporter, GFP-SKL (**Figure 1A**). GFP-SKL is a GFP with a C-terminal peroxisomal localization signal, which we have previously shown to be an accurate marker for peroxisomal dynamics (Chao *et al*. 2016; Wangler *et al*. 2017a). We anticipated that our screen could uncover three major categories of peroxisomal impact: Category A-an increase in the number of peroxisomes, Category Ban increase in the size of peroxisomes, and Category C-decrease or loss of peroxisomes or the marker (**Figure 1B**). We have previously observed examples of Category B in mutants with peroxisomal fission defects (Chao *et al*. 2016) and Category C in biogenesis defects (Wangler *et al*. 2017a) in *Drosophila*. We screened 215 lethal mutant lines from the collection that was mapped to a complementation group or to a gene (**Figure 1C, Supplemental Table 1**). We considered a hit to be positive when we could differentiate a clear difference in the mutant clone compared to the surrounding (heterozygous or homozygous wild-type) cells. In total we identified 37 alleles corresponding to 18 genes (**Table 1)**. For these hits, when possible we assayed 2 or more alleles per gene to confirm a change in GFP-SKL if possible. Some of these mutations led to an inconsistent increase of the peroxisome reporter amongst the fat body clones even within the same tissue, suggesting that perdurance of the wild type protein or other unmeasured factors may mask a change in peroxisomal dynamics in some clones. Interestingly, an overwhelming majority of the lines fell into Category A (**Table 1**). Although there are a few known peroxisomal related genes that are located on the Drosophila *X*-chromosome including *Pex5* (homolog of human *PEX5*) and *CG3415* (homolog of human *HSD17B4*) (Faust *et al*. 2012), we did not identify these in our screen. One explanation for this is that null alleles in a number of *Drosophila pex* genes have been shown to be viable (Nakayama *et al*. 2011; Faust *et al*. 2014; Wangler *et al*. 2017a), while the *X*-screen collection focused on recessive lethal mutations in essential genes. In contrast, none of the genes we isolated in this screen encode proteins that have been localized to peroxisome and are instead suggested to be involved in a wide variety of cellular processes (**Table 1**).

**Figure 1.**
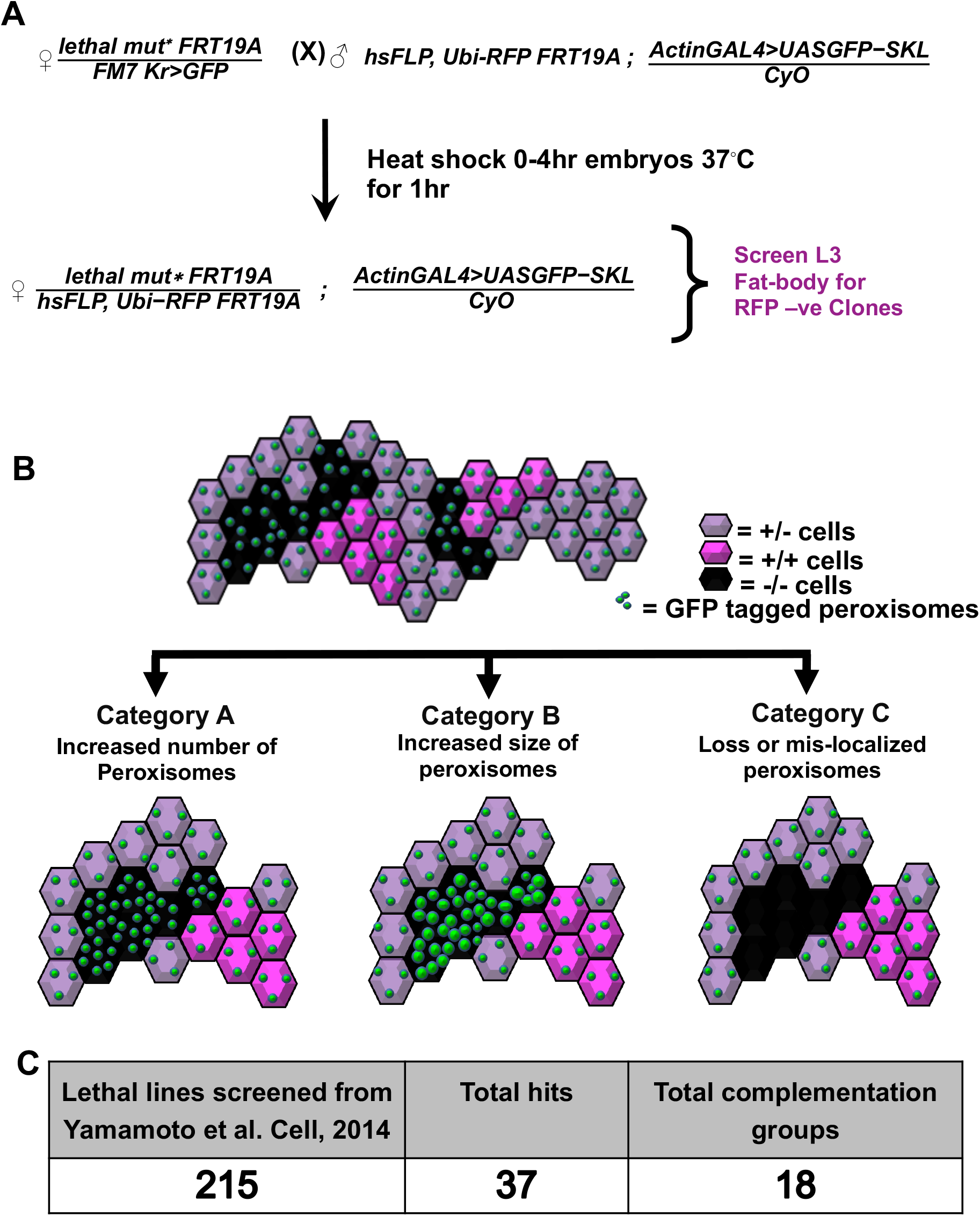
Overall study design and outcomes from the Drosophila X-Pex screen. A. Drosophila Cross Scheme showing in-detail. “<u>*lethal mut*</u>*” represents the different recessive lethal alleles used for the screen as listed in **Supplemental Table 1**. Males and females used for the experiment were crossed together at room temperature. Females were allowed to lay eggs for 4 hrs and the embryos were then heat-shocked at 37°C in a water-bath for 1hr and then kept at 25°C. The Fat bodies of the wondering third instar larvae were dissected, fixed and imaged by confocal microscopy. The homozygous mutant cells were identified through the absence of RFP (RFP-). B. Schematic representation of fat bodies expressing the GFP tagged peroxisome marker (GFP-SKL) with clones of mutant and wild-type cells. While homozygous mutant cells are marked by the absence of RFP, the sibling homozygous wild-type cell are marked by two doses of RFP (dark magenta). Heterozygous cells are marked with one dose of RFP (pale magenta) Category A represents an increase in peroxisomal numbers, Category B represents enlargement of peroxisomes, and Category C represents a loss of mislocalization of peroxisomal markers. C. Table representing the overall results from the screen. 215 total lines were screened, 37 total allele hits from 18 genes were identified.

In the majority of the lines screened, there was no difference between the appearance of the peroxisomal (GFP-SKL) marker in the mutant clone versus the surrounding sister cells, similar to the pattern seen in FRT19A controls (**Figure 2A-A”**). In one line, we observed strong enlargement and increased number of peroxisomes (**Figure 2B-B”**). This hit, *fs(1)h*, produced a Category A and B phenotype, and encodes a protein that has been reported to be involved in regulating proper expression of homeotic genes involved in pattern formation, such as *Ultrabithorax* (Florence and Faller 2008). However in other hits, including *Rbcn-3B, Coq8*, and *Usp16-45*, a Category A phenotype was observed with more GFP-SKL punctae in the clones (**Figure 2 C-E”**). Some of the hits from this screen have been studied in other biological contexts. *Rbcn-3B*, which produced a Category A phenotype, encodes a protein involved in Notch signaling during oogenesis through its role in endocytic trafficking and lysosomal function (Yan *et al*. 2009). *Coq8*, which produced a Category A phenotype, encodes a mitochondrial inner membrane protein that is predicted to be involved in electron transport (Zhu *et al*. 2017) In contrast, many of the genes we identified, including *Usp16-45* which is predicted to encode an ubiquitin specific protease, have not been extensively characterized *in vivo* and it is likely that these proteins also have additional functions as most genes are pleiotropic (Wangler *et al*. 2017b). These results indicate that screening for the GFP-SKL peroxisomal marker is an effective method to identify new genes that can impact peroxisome dynamics or morphology.

**Figure 2.**
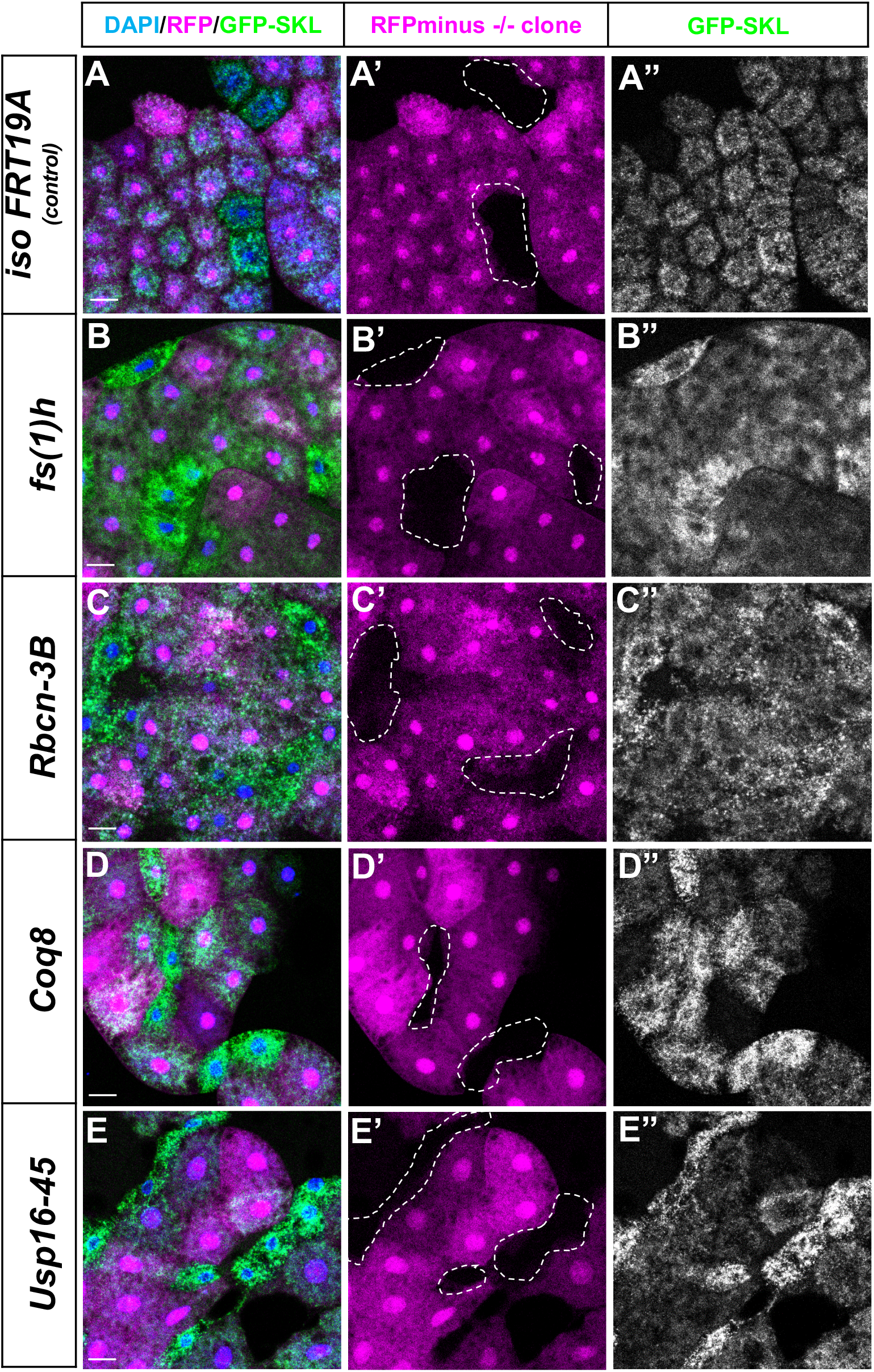
Peroxisomal morphological phenotypes observed *in-vivo*: Third instar fat body clones are shown in merge **DAPI (in blue) / RFP (magenta) / GFP-SKL (green)** in first from left column. Homozygous mutant cells that lack RFP are shown with dotted lines in the middle column and the right most column shows the same cells showing the **GFP-SKL** signal. The images in A-A”-*iso FRT19A* clones are clones of the non-mutagenized chromosome and serve as negative controls. Images of some of the strong hits are shown in panels B-E” as *fs(1)h* clones (from B-B”), *Rbcn-3B* clones (from C-C”), *Coq8 clones* (from D-D”) and *USP16-45* clones (from E-E”). Scale bars represent 50μm.

### Human homologs of genes identified in the fly peroxisome screen are candidates for human disease

In our previous studies from the *X*-screen collection we showed that the process of identification and screening for lethal mutations in *Drosophila* enriches for human Mendelian disease genes (Yamamoto *et al*. 2014). Indeed, our subsequent studies on this collection continued to yield novel human disease genes (*Deal and Yamamoto 2018*). We therefore wanted to assess this potential for genes from our peroxisome secondary screen within public human databases.

First, we determined the best human homolog of each fly gene. We examined each human homolog of the hits from our X-chromosome peroxisomal (X-Pex) screen in the Human Gene Nomenclature Orthology Prediction (HCOP) tool and the Drosophila RNAi Screening center Integrative Ortholog Prediction Tool (DIOPT) tool (Hu *et al*. 2011; Gray *et al*. 2015) (**Supplemental Table 2**). To access the latter, we utilized the MARRVEL tool, MARRVEL allows for simultaneous display of public human genomic data and model organism phenotypes and conservation (Wang et al. 2017). For the 18 fly genes that we considered hits, we found that sixteen of the eighteen had human homologs (88.9%). Thirteen out of sixteen fly genes had a single human homolog while remaining three of the genes had multiple human homologs (**Table 1**). We also wanted to know if any of the genes were already linked to a human single gene disorder. We examined the twenty human homologs from the X-Pex screen in a Mendelian disease using the Online Mendelian Inheritance in Man (OMIM) database (Amberger *et al*. 2015). Of the 23 human genes, nine are listed in relation to at least one Mendelian phenotype where the gene is causative for a described disease (**Supplemental Table 3**). Fourteen human genes have no known single gene disorder (**Table 2** and **Supplementary Table 3**). In our previous work in this collection, we observed that by screening for lethality, we enrich for essential genes in fly that are homologs of disease genes in humans (Wangler *et al*. 2015). We therefore hypothesized that these 14 human genes from our screen, not currently associated with human disease could be considered good candidates for undiagnosed cases.

**Table 2.** Human Gene Candidate Analysis. The human homologs of the X-Pex genes were examined for known Mendelian disease association (OMIM # entries) with genes that are not known to cause disease shown in red (Amberger *et al*. 2015). These are further sorted using data from the public human database gnomAD and the DOMINO scoring system for dominant disease. “pLI” score shows the probability (from 0-1) of a gene having intolerance to loss-of-function variation in the population of individuals represented in gnomAD data. “Missense z-score” show a z-score value for rates of missense variation in a gene. “pLI-o/e” is the observed / expected for loss-of-function variants in a gene, while “Mis-senze o/e” is a similar ratio for missense variants. The color code is red = o/e <0.2, orange= o/e <0.4, yellow = o/e <0.6, and gray represents o/e > =0.6. For DOMINO scores the code shows Red = “Very likely dominant (0.8-1)”, Orange = “Likely dominant (0.6-0.7)”, Indigo = “Either dominant or recessive (0.4-0.5)”, Blue = “Likely recessive (0.2-0.3), Turquoise = “Very Likely recessive (0-0.1)”

| Human Gene | Known Disease in OMIM | pLI | pLI o/e | Missense Z score | Missense Z score o/e | Domino |
| --- | --- | --- | --- | --- | --- | --- |
| **GSK3A** | None | 1 | 0 | 3.2 | 0.43 | 1 |
| **BRD4** | None | 1 | 0 | 3.74 | 0.63 | 0.997 |
| **UPF1** | None | 1 | 0.07 | 5.7 | 0.41 | 0.996 |
| **L1CAM** | # 304100, # 303350, # 307000 | 1 | 0.04 | 2.84 | 0.66 | n/a |
| **NFASC** | # 618356 | 1 | 0.12 | 2.59 | 0.74 | 0.871 |
| **BRD3** | None | 0.98 | 0.14 | 3.76 | 0.64 | 0.893 |
| **GSK3B** | None | 0.96 | 0.14 | 2.91 | 0.48 | 1 |
| **SMAD3** | # 613795 | 0.84 | 0.17 | 3.47 | 0.39 | 0.999 |
| **UPF2** | None | 1 | 0.03 | 3.3 | 0.65 | 0.723 |
| **WDR7** | None | 1 | 0.10 | 2.63 | 0.75 | 0.636 |
| **BRD2** | None | 1 | 0.08 | 0.5 | 0.93 | 0.474 |
| **MTPAP** | # 613672 | 1 | 0.04 | 1.05 | 0.84 | 0.242 |
| **PTAR1** | None | 0.85 | 0.15 | 1.51 | 0.71 | 0.275 |
| **NRCAM** | None | 0.18 | 0.24 | 2.05 | 0.79 | 0.191 |
| **FH** | # 606812 - # 150800 | 0.09 | 0.28 | 1.39 | 0.77 | 0.371 |
| **HINFP** | None | 0.03 | 0.29 | 1.73 | 0.73 | 0.719 |
| **PI4KA** | #616531 | 0 | 0.36 | 3.53 | 0.72 | 0.589 |
| **BRDT** | # 617644 | 0 | 0.46 | 0.47 | 0.94 | 0.209 |
| **CHL1** | None | 0 | 0.47 | -1.92 | 1.21 | 0.152 |
| **USP45** | None | 0 | 0.74 | 0.75 | 0.90 | 0.168 |
| **COQ7** | # 616733 | 0 | 0.88 | -0.42 | 1.10 | 0.091 |
| **RFT1** | #612015 | 0 | 0.77 | 0.97 | 0.84 | 0.063 |
| **COQ8B**<br>(a.k.a. ADCK4) | # 615573 | 0 | 0.78 | 0.89 | 0.87 | 0.056 |

In order to explore evidence for this we examined public human genomic databases, we were looking for evidence of selective constraint in all 23 human genes identified by our screen. We examined the gnomAD database which is a large genome and exome aggregation largely selected for healthy or adult-onset disease cases (Lek *et al*. 2016). For each gene we examined the constraint metrics or evidence that damaging variants in the gene are absent from these “control” individuals (**Table 2, Supplemental Table 3**). Twelve of the 23 genes had “observed over expected” (o/e) numbers for loss-of-function variants at 0.2 (20%) or less indicating fewer loss-of function alleles than anticipated by chance. This data suggests that there may be some selection against loss of function alleles for approximately half of the genes possibly due to a haploinsufficient mechanism. We compared the characteristics of these genes to two other sets of genes, first we compared the X-Pex gene set to all the other homologs of the genes that we screened. We also compared to a group of 25 well known human peroxisomal disease genes that encode proteins in the peroxisome biogenesis machinery and enzymes involved in very-long-chain fatty acid oxidation, plasmalogen synthesis and reactive oxygen species (**Supplemental Table 4**). Comparing these three gene sets we found that the human homologs of the essential fly genes had significantly higher probability of loss of function intolerance (**Figure 3A-B**), and higher missense constraint (**Figure 3C-D**) in the human databases compared to the known peroxisomal disease genes. This was consistent across both genes that were positive in the peroxisome secondary screen as well as negative. These intolerance scores apply more to dominant disorders than autosomal recessive. Consistent with that, the majority of the known peroxisomal genes underly recessive disease (**Supplemental Table 4**). To date, very few dominant peroxisomal diseases have been identified. We therefore hypothesized that the selection of lethals in our original fly screen pointed us to a set of human genes that are more likely to underly dominant disease. The constraint metrics of the gnomAD dataset are indeed most valuable for showing selective constraint for heterozygous alleles, such as *de novo* mutations or dominant inherited disorders, particularly with early onset or an impact on reproduction (Lek *et al*. 2016).

**Figure 3.**
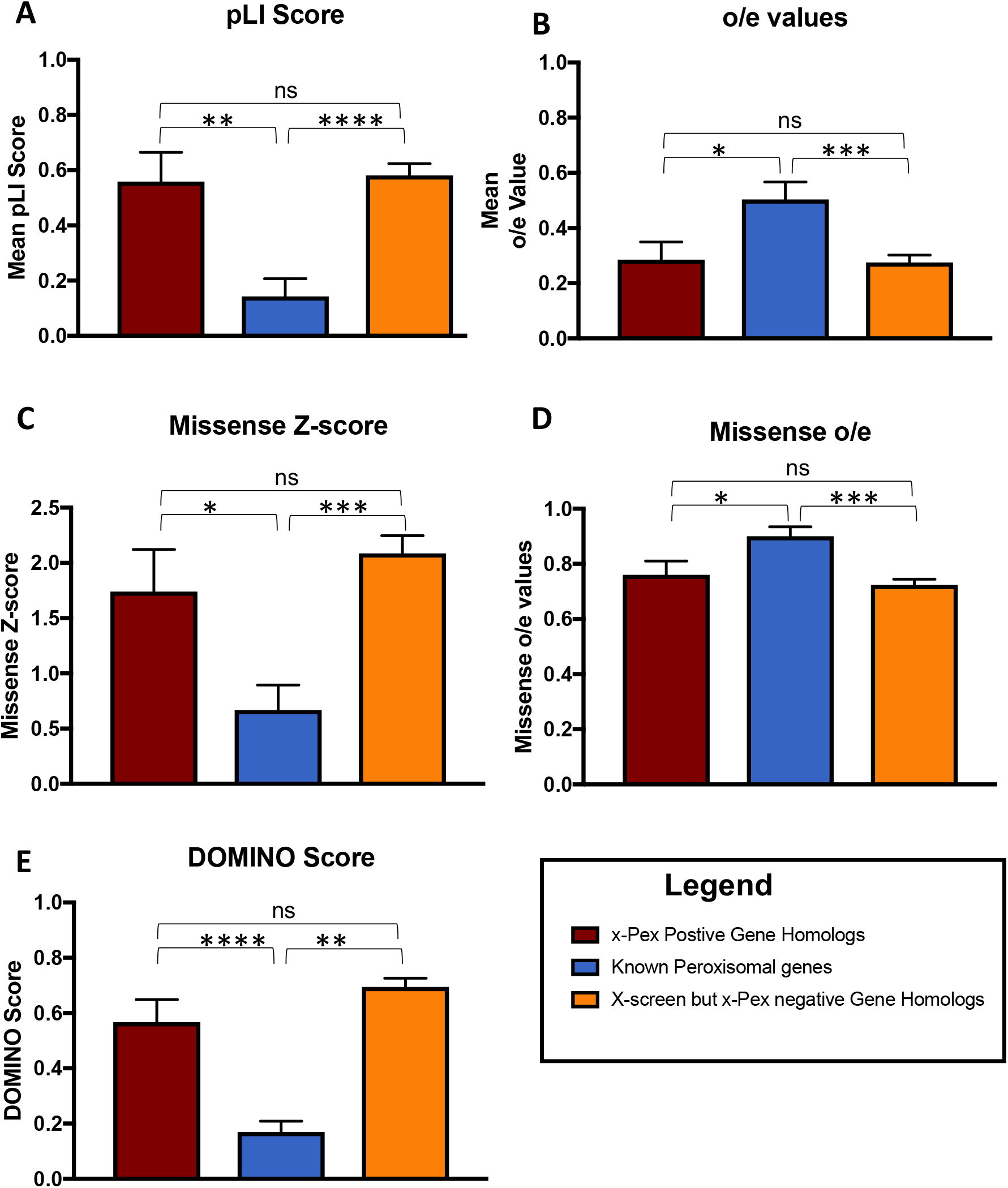
Comparison of known human peroxisomal disease genes to the new X-Pex candidates. A. The Probability of Loss of Function intolerance score (pLi) calculated from public human data from the gnomAD database (Lek *et al*. 2016). The X-Pex genes displayed a mean pLi score of 0.55 ± 0.11, n=20, while the known peroxisomal disease genes had a mean pLi of 0.14 ± 0.06, n=25, which was statistically significant (p=0.0016) **. B. The observed over expected (o/e) loss of function scores calculated from public human data from the gnomAD database. The X-Pex genes had a mean o/e score of 0.29 ± 0.06, n=20, while the known peroxisomal disease genes had an o/e score of 0.50 ± 0.06, n=25, which was statistically significant (p= 0.0218)*. C. The missense constrain z-score calculated from public human data from the gnomAD database. The X-Pex genes had mean missense constrain z-scores of 2.16 ± 0.34, n=20, while the known peroxisomal genes had z-scores of 0.67± 0.23, n=25, which was statistically significant (p=0.0005)***. D. Missense constraint o/e scores calculated from public human data from the gnomAD database. The X-Pex genes had a mean o/e for missense variants of 0.73 ± 0.04, n=20, compared to the known peroxisomal disease genes o/e score of 0.90 ± 0.03, n=25, also statistically significant (p=0.0025)**. E. DOMINO scores calculated for the gene sets. The X-Pex gene set had a DOMINO score of 0.53 ± 0.08, n=20, while the known peroxisomal disease genes had a mean DOMINO score of 0.17 ± 0.04, n=24, and the difference was statistically significant (p<0.0001)***.

In order to test this, we used the DOMINO tool to assess the probability of dominant disease for each gene (Quinodoz *et al*. 2017). The DOMINO score, indicating the likelihood of dominant disorders also differed between the X-Pex gene set and the known peroxisomal disease gene set (**Figure 3E**). The X-Pex gene set had a higher DOMINO score, while the known peroxisomal disease genes had lower DOMINO scores, thus more likelihood of relating to recessive disease and the difference was statistically significant (p<0.0001). As noted, for the known peroxisomal genes this is indeed the case, as 22 of the 25 genes are disease genes for autosomal recessive disorders (**Supplemental Table 4**).

With this data we predict that some X-Pex genes could underlie dominant phenotypes, we sought evidence for dominant phenotypes related to alleles in the set of X-Pex genes using public databases of *de novo* events from individuals with disease and controls (Turner *et al*. 2017). This dataset primarily focuses on neurodevelopmental phenotypes and the *de novo* events from diverse cohorts. Strikingly, we observed suggestive results for six genes from the X-Pex gene set with high DOMINO scores (*GSK3A, BRD4, UPF1, BRD3, GSK3B* and *SMAD3*) and at least one individual with developmental delay, Autism, or congenital heart disease. These cases are not definitively linked to these loci and are noted to have a missense *de novo* event. Interestingly no missense *de novo* events are observed in control individuals for these genes (**Supplemental Table 5**).

Whether these genes are ultimately good candidates remains to be explored in undiagnosed cases as these database searches do not definitely link the specific gene with disease. It is also not known whether the peroxisomal phenotype that was observed in our screen would be conserved in humans. Peroxisomes are not routinely examined in clinical samples, so this data provides a key starting point for these genes of interest. Even a secondary impact on peroxisomes without direct interaction could aid in exploring these candidate genes further.

Taken together we propose the X-Pex provides a good list of candidate genes, in particular, *de novo* events in these genes from patients with neurodevelopmental phenotypes should be explored. Considering that peroxisomal disease classically relates to autosomal recessive conditions, the X-Pex gene list may provide an entry point to study the role of *de novo* events in genes that impact peroxisomes that have been missed in previous screens. This study therefore provides additional support for the use of forward genetic screens in model organisms in the study and identification of human disease genes (Yamamoto *et al*. 2014; Wangler *et al*. 2015).

## Acknowledgments

The authors thank the Kyoto and Bloomington *Drosophila* Stock Centers for maintaining and publically distributing the X-chromosome recessive lethal stocks used in this study. S.Y. and M.F.W thank the Junior Faculty Seed Funds from the Department of Molecular and Human Genetics at Baylor College of Medicine and Global Foundation for Peroxisomal Disorders, RhizoKids and Wynne Mateffy Research Foundation to support this research.

